# The role of SRSF3 splicing factor in generating circular RNAs

**DOI:** 10.1101/799700

**Authors:** Ammar S. Naqvi, Mukta Asnani, Kathryn L. Black, Katharina E. Hayer, Deanne Taylor, Andrei Thomas-Tikhonenko

## Abstract

Circular RNAs (circRNAs) represent a novel class of non-coding RNAs that are emerging as potentially important regulators of gene expression. circRNAs are typically generated from host gene transcripts through a non-canonical back-splicing mechanism, whose regulation is still not well understood. To explore regulation of circRNAs in cancer, we generated sequence data from RNase R-resistant transcripts in human p493-6 B-lymphoid cells and identified thousands of novel as well as previously identified circRNAs. Approximately 40% of expressed genes generated a circRNA, with half of them generating multiple isoforms, suggesting the involvement of alternative back-splicing and regulatory RNA-binding proteins (RBPs). We observed that genes generating circRNAs with back-spliced exonic junctions were enriched for RBP recognition motifs, including multiple splicing factors, most notably SRSF3, a splicing factor known to promote exon inclusion. To test the role of SRSF3 role in circRNA production, we performed traditional RNA-seq in p493-6 B-lymphoid cells with and without SRSF3 knockdown, and identified 926 mRNA transcripts, whose canonical splicing was affected by SRSF3. We found that a subset (205) of these SRSF3 targets served as host transcripts for circRNA, suggesting that SRSF3 may regulate exon circularization. Since this splicing factor is deregulated in hematologic malignancies, we hypothesize that SRSF3-dependent circRNAs, similar to their mRNA counterparts, might contribute to the pathogenesis of lymphomas and leukemias.

## Introduction

CircRNAs are a relatively new class of non-coding RNAs with covalently closed 5’ and 3’ ends. The vast majority of circRNAs originate from exons, although there have been studies showing that they are produced from a variety of regions in the genome, including introns, exons, and UTR regions. The cytoplasm is host to the majority of circRNAs (Kristensen et al., 2018), but there is recent evidence of certain circRNAs enriched in the nucleus and controlled or modulated by sequence length (Huang et al., 2018). Interestingly, most circRNAs are not more abundant when compared to corresponding mRNA transcripts, but circRNAs exhibit greater half-lives demonstrating their increased stability (Kristensen et al., 2018).

The biological role of circRNAs still remains fairly elusive, but with increasing evidence towards performing multiple functions. The most well cited functionality involves a microRNA (miR) “sponge” role, where circRNAs effectively control proteins and target mRNAs by binding to and reducing the availability of miRs. CircRNAs may have functional relationships with genes and miRs known to have association with cancer pathogenesis. One notable example is the sponge activity against miR-7 which is a known modulator of several oncogenes. The circRNA ciRS7 (also known as Cdr1as) contains approximately 70 conserved binding sites for miR-7 and is widely associated with Argonaute (AGO) proteins. Overexpression of ciRS7 results in increased miR-7 suppression leading to a concomitant increase in miR-7’s target levels (Hansen et al., 2013; Memczak et al., 2013). Another circRNA that has been shown to target miR-7 is circHIPK3. Knockdown of circHIPK3 markedly inhibits in vitro colorectal cancer cells proliferation, migration, invasion, and induces apoptosis. It also is observed to be upregulated in colorectal cancer tissues and cell lines (Zeng et al., 2018). Similarly, circRNAs could control steady state levels of proteins encoded by their host genes. For instance, PABPN1-derived circRNA has the ability to control steady state levels of the protein PABN1 through sponge activity against HuR, a RNA-binding protein that assists in translation (Abdelmohsen et al., 2017). In another case, circRNAs could contain binding sites for multiple proteins, acting as scaffolds by bringing two proteins together and mediating an interaction. In one study, circFOXO3 mediates the interaction of MDM2 and key tumor suppressor p53, implicating it in growth arrest, senescence, and apoptosis (Du et al., 2017). Given some of these characterized and putative functions, circRNAs are emerging as important players in the cancer field (Abdelmohsen et al., 2017; Hansen et al., 2013; Meng et al., 2017; Patop and Kadener, 2018; Zhang et al., 2017).

Although it is well established that circRNAs are produced from a distinct back-splicing process (Wang and Wang, 2015), the details of circRNA biogenesis still remain very much elusive. There have been thousands of predicted circRNAs with multiple databases (Ghosal et al., 2013; Glazar et al., 2014; Xia et al., 2018; Zhang et al., 2016) housing these candidates, but how circRNAs are generated from host genes is still being elucidated. Some reports suggest that some circRNAs are back-spliced when two exons are brought close to each other through the mediation of Alu elements contained in the flanking introns (Liang and Wilusz, 2014). However, some studies point out that back-splicing of exons occurs when RNA binding proteins (RBPs) that bind to the flanking introns of the back-spliced exons, dimerize, bringing exons closer to each other, as the case for the RBP QKI (Conn et al., 2015).

Here, we describe our mechanistic studies performed on EBV-transduced P493-6 cells, which is frequently used as a model cell line to study Burkitt lymphoma (Pajic et al., 2000). Our results suggest that in P493-6 a combination of these mechanisms mediate the back-splicing of circRNAs. It also suggests that splicing factors, such as SRSF3, could be key to the generation of this class of non-coding RNAs. Moreover, the dysregulation of some of these splicing factors could control the expression of at least a subset of circRNAs that would then entail important functional consequences that could contribute to neoplastic transformation of B-cells. This has already been shown by us in the context of aberrant splicing of SRSF3-dependent mRNA transcript encoding CD19 (Asnani, 2019; Black et al., 2018; Sotillo et al., 2015), but here we provide increasing evidence that gene dysregulation could also occur via circRNAs.

## Results

### circular RNAs expressed in P493-6 cells

In order to identify circRNAs, we followed standard protocols for circRNA-seq (see Methods), which involves a ribo-depletion step (Fig. 1A), followed by RNAse-R treated RNA-seq. We designed a pipeline to identify bona-fide exonic circRNAs (Fig. 1B), where we combined readily available tools such as mapSplice2 and integrated our customized filters to take advantage of replicates, known annotations and conservation. Although there are multiple tools available for circRNA detection, we chose mapSplice2, which was the most conservative and accurate tool, when benchmarked against four other popular methods (Hansen et al., 2016). Using p439-6 cells, we were able to identify 957 total host genes or transcripts that corresponded to 1324 circRNAs. In order to independently confirm our results, we were able to rediscover our circRNA candidates by overlapping our host transcripts and circRNAs with MC116 cells, which are a similar cell line from a previous study (Fig. 1D) (Tagawa et al., 2018). The non-overlapping circRNAs are most likely due to different techniques and algorithms used for circRNA detections, as we used a more conservative method. Interestingly, the other non-overlapping circRNAs were previously not reported, suggesting a subset of novel circRNAs in our cell system.

**Figure 1.**
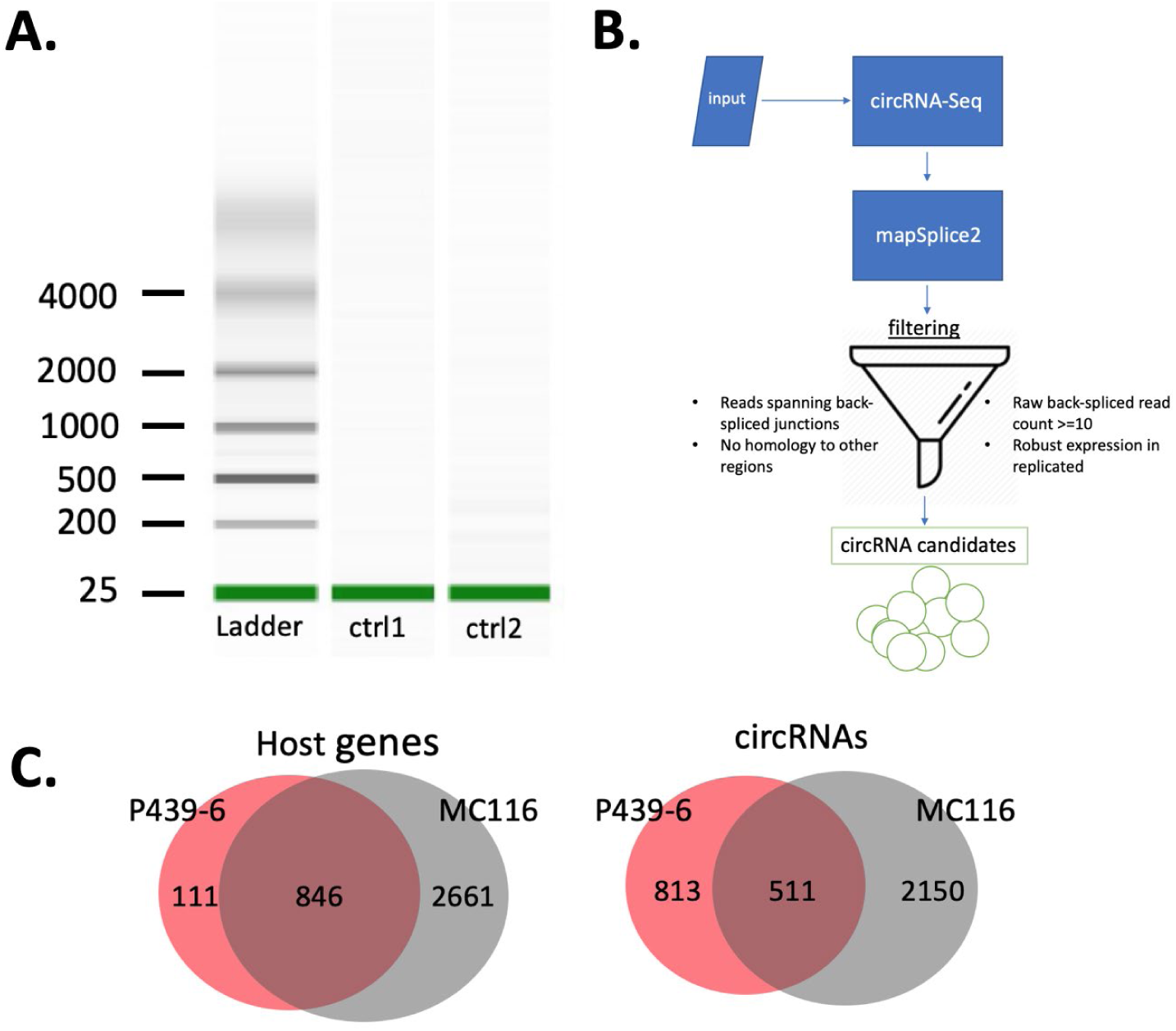
Total circRNAs identified. **(A)** Blot showing successful Ribo-depletion of total RNA in p493-6 cells. Controls (ctrl1, ctrl2) show no ribosomal or artifactual RNA. **(B)** Computational pipeline and workflow of circRNA identification where input is an RNA-seq data file. **(C)** Venn diagrams of host genes and circRNAs found in P439-6 and MC116 cells lines, discussed in text.

Additionally, we computed the number of circRNA isoforms per host gene (Fig. 2A) and we observed almost one-third of these transcripts producing multiple isoforms, suggesting the propensity of circRNAs to be alternatively back-spliced, confirming past findings (Glažar et al., 2014). For example, we discovered a single circRNA species processed from a single transcript representing a one-to-one relationship, as in the case for B2M, where exons one and two are back-spliced into a circRNA (Fig. 2B). Additionally, we observed an alternatively back-spliced circRNA was processed from the host transcript FBXW7. Specifically, we observed that there was an alternative exon assembly that resulted in three different circRNA isoforms consisting of four distinct exons, processed from both the 5’ end and 3’ ends of the host gene (Fig. 2C).

**Figure 2:**
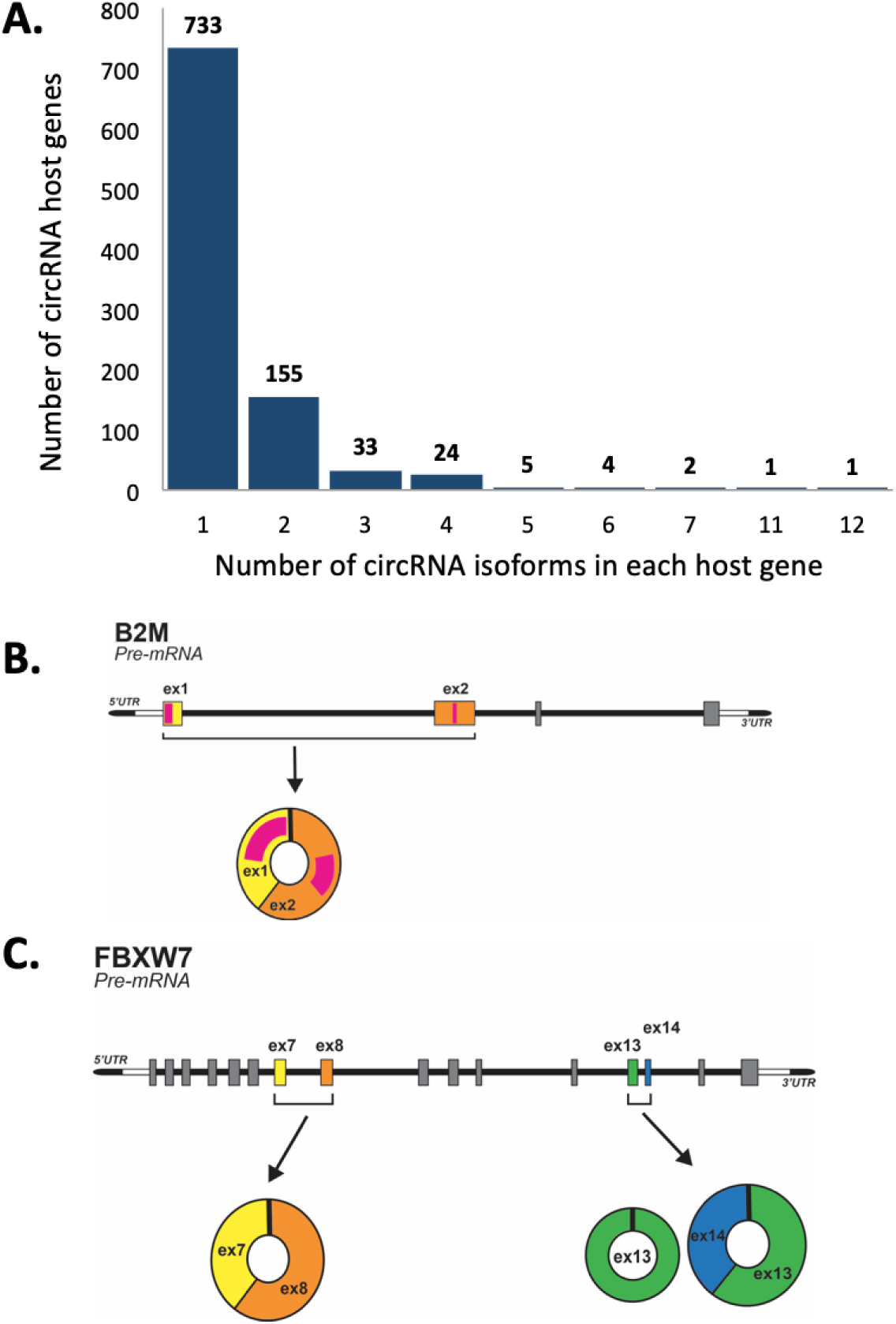
Isoforms of circRNAs **(A)** Histogram representing the number of isoforms generated by host genes; for example there are 733 host genes each producing one circRNA isoform. corresponding to one gene. **(B)** One circRNA isoform is observed from one host gene. **(C)** Example of multiple circRNA isoforms observed from one host gene, here 3 isoforms are produced from FBXW7.

### Alu vs non-alu circRNAs

In order to test previously held models of circRNA biogenesis, we sought to investigate the proposed relationship between Alu elements and back-splicing. (Liang and Wilusz, 2014). Thus, we first searched for known Alu elements in regions flanking the two exons that made up the back-spliced portion of each circRNA in our candidate circRNA dataset. We found that 840 circRNAs contained Alu elements, with only a subset of them being consistent with the model (Fig. 3A), while 484 circRNAs did not contain flanking Alu elements. For example, circRNAs associated with the transcript GNBL contained Alu elements, but only one of these circRNA isoforms followed the previous models of flanking Alu elements mediating the back-splicing of exons to form the back-spliced junction (Fig. 3B).

**Figure 3:**
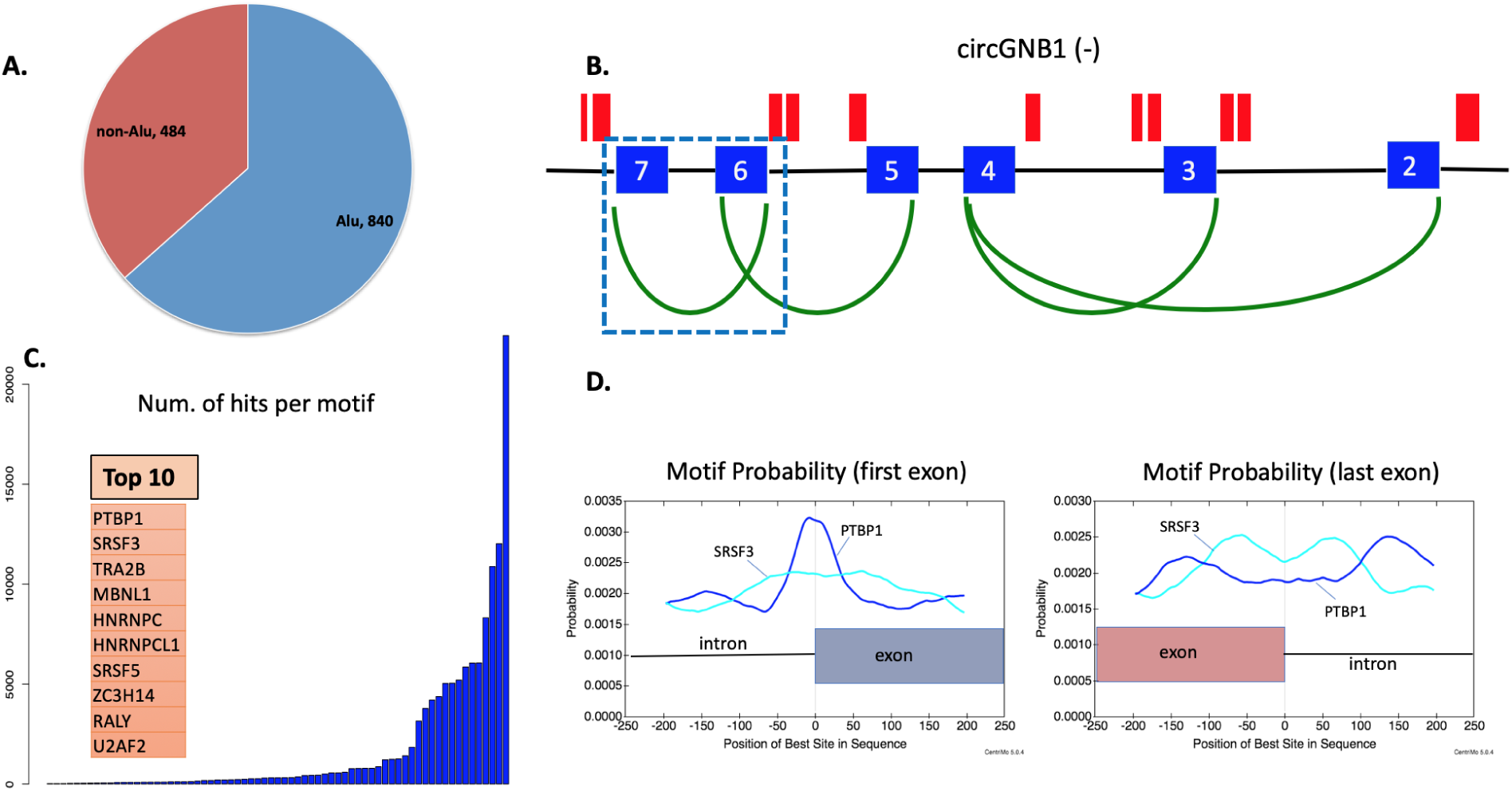
Presence of Alu elements and RNA-binding proteins in circRNA sequences. **(A)** Pie chart indicating the total number of known Alu elements in flanking intron sequences of back-spliced exons from circRNA candidate sequences. **(B)** Example of highly expressive circGNB1 back-splicing (green arcs) with indicated Alu regions in red. Exons 7 and 6 (squared) are the only back-splicing case following the Alu-based model for circRNA biogenesis. **(C)** Number of hits per each known RNA binding motif in sequences of back-spliced exons and flanking introns (+/-250nts). **(D)** Positional probability of SRSF3 and PTBP1 motif occurrences in back-spliced exons and corresponding flanking intron sequences.

### SRSF3’s role in circRNA processing

In order to complement the above analysis, we tested the alternative hypotheses related to the involvement of RBPs and their role in circRNA biogenesis. Using Multiple EM for Motif Elicitation (MEME) (Bailey et al., 2006), we searched for known RNA-binding motifs partially spanning the 5’/3’ flanking introns and exons (+/− 250nts), including the intron-exon boundaries of the first and last exons. We were able to detect multiple binding motifs, with our top hits consisting of known splicing factors, including PTBP1 and SRSF3 (Fig 3C). Interestingly, we also observed preferential motif positions in these sequences, specifically for SRSF3 and PTBP1 (Fig. 3D). Our results indicate that these regions may be bound by multiple RBPs at distinct nucleotide positions revealing a positional binding bias, as they do in mRNA processing. Since the majority of circRNAs are exonic and SRSF3 is commonly known to promote exon inclusion, we focused on this particular splice factor.

To expand on this idea, we sought to identify alternative splicing targets of SRSF3. To model downregulation of SRSF3 and to identify affected mRNA and circRNA transcripts, we knocked down SRSF3 mRNA with siRNA in the P493-6 cell line. Both the non-targeting scrambled control and the SRSF3-targeting reagent were Smart Pools of four siRNAs, designed to target distinct sites within the specific gene of interest. This approach allowed us to reduce the reliance on and the concentration of each individual siRNA and average out their off-target effects, as documented by (Parsons et al., 2009). We performed this experiment at three different time intervals, including 24, 48 and 72 hours. Knockdown of SRSF3 was confirmed by RT-qPCR (Figure 4A) and Western blotting (Figure 4B). We then performed RNA-seq on the 72hr RNA population, when the best SRSF3 protein down-regulation took place. Following RNA-Seq and Modeling Alternative Junction Inclusion Quantification (MAJIQ) (Vaquero-Garcia et al., 2016) analysis, we found 926 mRNAs that underwent differential splicing between the control vs SRSF3 knock-down conditions, identifying targets of SRSF3. Of these, more than a fifth (or 205) also corresponded to circRNA candidates that we identified, including those that had enrichment for SRSF3 motifs (Fig 4). Thus, our data suggest that the dysregulation of SRSF3 does not only influence the splicing of mRNA targets, but also affects the back-splicing of circRNAs.

**Figure 4.**
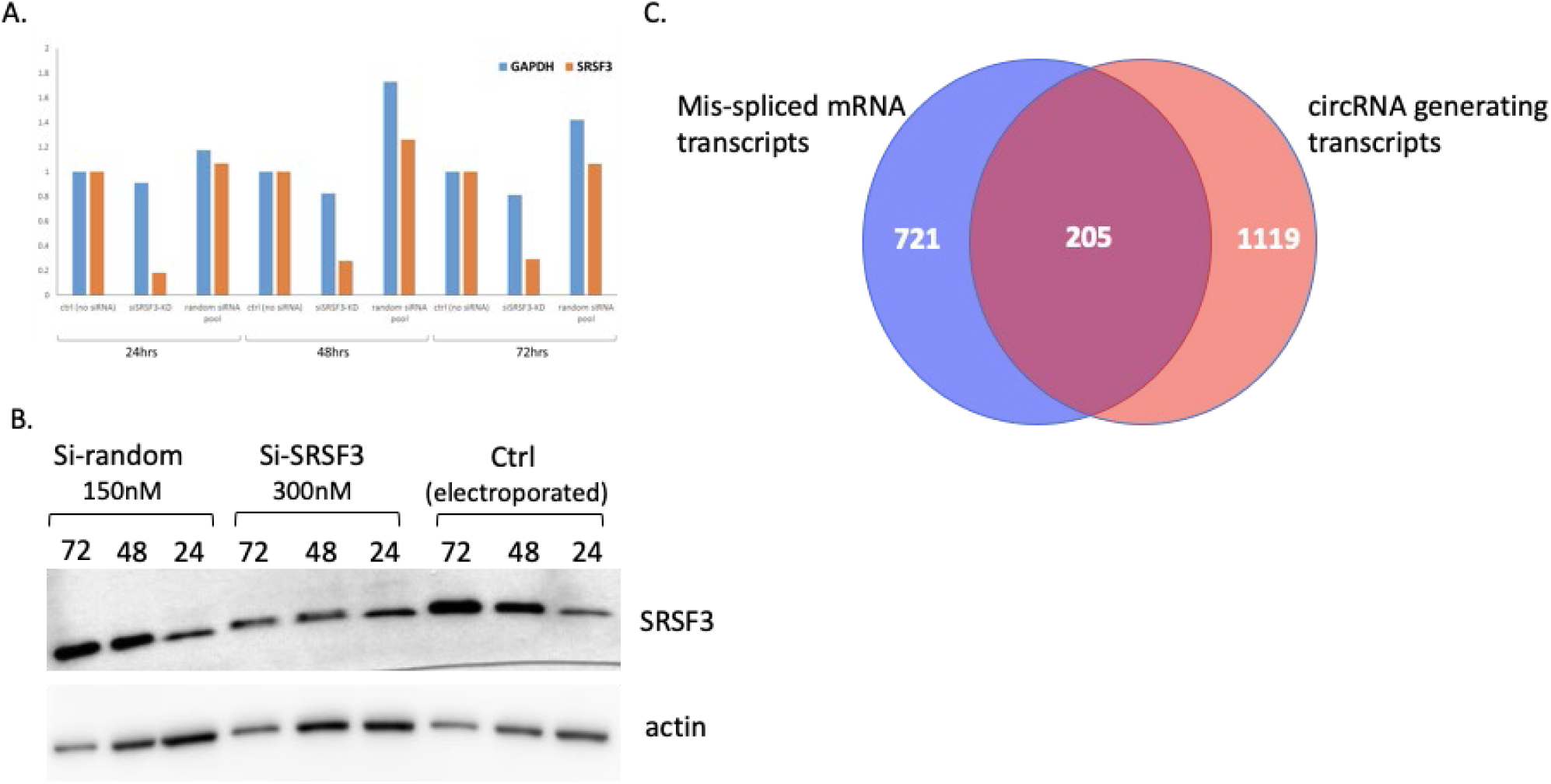
SRSF3 Knock-down in p493-6 cells **(A)** qRT-PCR validation of control vs knock-down in 24, 48 and 72 hours. **(B)** Western blot of protein levels upon knocking down SRSF3 in 24, 48, and 72 hours. (**C)** Venn diagram of mis-spliced mRNA transcripts upon SRSF3 knockdown and transcripts generating circRNAs.

## Discussion

Our findings reveal that SRSF3 may be an important player in circRNA biogenesis. As we performed circRNA-seq on the P493-6 cells, we found that there were many circRNA candidates that were enriched for motifs of known splicing factors. This corroborates the proposed model of RBP mediated back-splicing. Furthermore, our findings also suggest alternative back-splicing within these candidates. We also found evidence for SRSF3-dependent transcripts to produce circRNAs, indicating that SRSF3 not only controls the splicing of mRNAs, but also circRNAs. Taken together, this positions us to study the relationship of the role of exon skipping/inclusion and circRNA back-splicing. For example, we will determine whether an increase in exon skipping entail increased circRNA processing and vice versa.

Interestingly, we have also shown that SRSF3 is mis-spliced in primary leukemias with increased inclusion of poison exons (Anko et al., 2012; Black et al., 2018). It has also been shown that SRSF3 binds to TP53 mRNA, resulting in an alternatively spliced isoform of p53 that promotes cellular senescence(Tang et al., 2013). The perturbation of this interaction is thought to lead to profound functional consequences in cancer. Expanding this hypothesis, we could postulate that the dysregulation of SRSF3 (and other splicing factors) could also alter a subset of circRNAs that then would have an effect on its downstream targets. In other words, the dual functions of modulating canonical and non-canonical splicing (back-splicing) by SRSF3 and other RBPs will simultaneously affect both mRNAs and circRNAs. This will presumably affect circRNA sponge activity (Hansen et al., 2013; Memczak et al., 2013) disrupting the steady state levels of proteins and mRNAs in premalignant cells.

## Materials and methods

### Conventional RNA-seq

RNA was isolated using Qiagen RNeasy Mini Kit. RNA integrity and concentration were determined using Eukaryote Total RNA Nano assay on BioAnalyzer. RNA-seq was performed on 2ug of total RNA according to the GeneWiz Illumina Hi-seq protocol for ribo-depletion samples (2 × 150 bp pair-end sequencing, 350M raw reads per lane).

### RNAse-R treated RNA-seq of p493-6 cells

To enrich for circRNAs, the total RNA pool was used for an RNase R (Epicentre, Illumina, San Diego, CA, USA) digestion (16 units RNase R, 15 minutes at 37uC) to deplete linear and enrich circular RNAs. A 10-fold volume or 500 ml of QIAzol Lysis Reagent and 3-fold volume of chloroform was added to the RNase R reaction. After mixture and phase separation, the aqueous phase was extracted with 600 ml chloroform. 1 ml glycogen was added to the aqueous phase and nucleic acids precipitated with a 1-fold volume of isopropanol. The pellet was then washed with 70% ethanol and resuspended in water. A second round of RNase R digestion, QIAzol/chloroform extraction and isopropanol precipitation followed. RNA-seq was performed on 20-170 ng of total RNA according to the GeneWiz Illumina Hi-seq protocol for ribo-depletion samples (2 × 150 bp paired-end sequencing, 350M raw reads per lane).

### circRNA identification

Fastq files of RNA-seq obtained from GeneWiz were mapped using mapsplice2 (Wang et al., 2010). Paired-end reads that mapped and spanned the back-spliced junctions were used for downstream identifications, quantifications and differential expression analyses. Additionally, mapped reads were discarded if: 1. there were < 10 raw reads in any of the replicates that spanned the back-spliced exon junctions, and 2. reads aligned somewhere else in the genome.

### Motif detection and positional probability of SRSF3 and PTBP1 motif occurrences

The motif detection pipeline utilized RBPMap (v. 1.1) with a medium stringency level (*P*-value ≤ 0.005) and conservation filter (Paz et al., 2014). The analysis was performed on sequences 250 +/− nts from the intron-exon and exon-intron boundaries of the back-spliced exons or the first and last exons, respectively. All known RNA-binding motifs were identified and average z-scores and *P*-values for all significant hits were calculated. Probabilities of known SRSF3 and PTBP1 motif occurrences in these circRNA sequences were calculated using Find Individual Motif Occurrence or FIMO program (-v 5.05) (PMC3065696).

### RNA-Seq alignment, quantification and differential expression

Fastq files of RNA-seq obtained from GeneWiz were mapped using STAR aligner. STAR was run with the option ‘alignSJoverhangMin 8′. STAR genome reference were based on the hg19 build. Alignments were then quantified for each mRNA transcript using HTSeq with the Ensembl-based GFF file and with the option ‘-m intersection-strict’. Normalization of the raw reads was performed using the trimmed mean of M-values (TMM). Differential expression of wild-type and knock-down or pro-B and B-ALL RNA-Seq datasets were assessed based on a model using the negative binomial distribution, a method employed by the R package DESeq2. Those differential genes that had a P-value of < 0.05 were deemed as significantly up or down-regulated.

### Splicing analysis

In order to detect local splicing variations (LSVs), the MAJIQ tool (-v 2.03) was used. MAJIQ was run on the Ensembl-based GFF annotations, disallowing de novo calls. LSVs that had at least a 10% change at a 95% confidence interval between two given conditions were chosen for further analysis. The 95% confidence interval in this context is defined as follows: given observed reads in both experiments there is a 95% posterior probability, according to the MAJIQ model, that there is a change of 10% or more in PSI.

